# Investigation of nano- and micro-domains formed by ceramide-1-phosphate in freestanding lipid bilayers

**DOI:** 10.1101/2022.06.01.494282

**Authors:** Dominik Drabik, Aleksander Czogalla

## Abstract

Biological membranes are known for their complex nature and formation of domains is crucial for proper execution of multiple cellular processes. Such domains, mostly due to their nanoscale character, are rarely studied since implementation of techniques for their quantitative investigation is difficult. In this article we introduce spot-variation z-scan FCS implemented for the first time for artificial lipid vesicles of defined lipid composition. We used this approach to investigate behaviour of different species of ceramide-1-phosphate within membranes. We were able to provide quantitative description of lipid domains generated by ceramide-1-phosphate in nano and microscale. Aligning these results with in silico studies helped us to validate the approach and draw conclusions on complex behaviour of signalling lipids within biological membranes.

---

Biological membranes are recognized as complex structures of heterogeneous nature. Existence of lateral domains are one of the hallmarks of organellar and cellular membranes. Plasma membranes consist of many coexisting dynamic domains, that vary in composition and size (from nano to micrometers in diameter). Such liquid orderered (L_o_)/liquid disordered (L_d_) domain coexistence can also be reconstituted in simple model membrane systems [1, 2]. In living cell systems, raft domains, which are more ordered in their nature, exist as small (<100nm), highly dynamic protein-lipid entities induced by their interactions. Upon signaling, raft nano domains may coalesce into larger micrometer domains (>200 nm) and act as sorting platforms, enabling the selective association of specific molecules to subsequently fulfill their role in numerous cellular processes [3, 4]. In artificial lipid membrane systems, such as giant unilamellar vesicles (GUVs), such micro domains are usually formed in time regime of tens of μs. This makes GUVs ideal system for domain visualization and study.

One of techniques that allow investigation of nano-domains is spot-variation fluorescence correlation spectroscopy (svFCS). It is one of the most promising single-molecule approach to study membrane domain related phenomena, as it was already successfully implemented to study deeply the dynamics of cellular plasma membrane organization [5, 6]. In this technique the diffusion time against squared waist of focal volume is plotted [7, 8]. This allows to determine two observables - effective diffusion coefficient (D_eff_) and the intercept of the diffusion time axis (t_0_) that is indicative of potential confinement of studied molecules within membrane domains of various character. It was established that higher values of t_0_ corresponds to dynamic clustering, which influence the free movement. On the other hand, negative t_0_ is supposed to be related to so called meshwork barriers. In this paper, we established a new protocol based on svFCS and GUVs enabling studies of diffusivity within freestanding lipid bilayers. We report, for the first time to our knowledge, successful implementation of svFCS methodology to study diffusion coefficient determination and nano-domain presence in freestanding lipid membranes. We combined this technique with z-scan FCS approach [9, 10], which opens new perspectives for calibration-free methodology applicable for lipid vesicles. The obtained results were compared with classical diffusion coefficients calculated from molecular dynamics simulations.

We’ve selected membranes containing important signaling lipid, ceramide-1-phoshopate (C1P), that regulates cell proliferation, apoptosis and acts as a mediator of inflammatory responses [11, 12]. This sphingoid analogue of phosphatidic acid (PA) also has the ability to induce curvature of lipid bilayers. Since C1P is a signaling molecule, its phase behavior and ability to form domains is of particular interest. However, only a few literature reports exist in this regard, which are self-conflicting in conclusions [13–17]. In addition to study nano-domains within C1P enriched membranes, additional insight into domain-inducing nature of the lipid was obtained via microscopical analysis of micro domains.

In order to establish the protocol for svFCS on GUVs, the initial measurements were performed on single component POPC vesicles with NBD-Chol fluorophore. The initial results showed considerable deviations between diffusion times for given vesicle population under constant waist size. This led us to a conclusion that the issue might exist due to imperfect positioning of the point spread function relative to the plane of the membrane, while considering only the highest fluorescence intensity. To address this issue, we decided to adapt z-scan approach. More specifically, the position in Z-axis of strongest intensity as well as two positions above and below this position are selected. Then obtained diffusion times are plotted as a function of z-position and fitted with parabolic function to get minimum diffusion time for given vesicle which should be considered as the most reliable (as shown in Figure 1.B). After establishing the value for at least 5 vesicles, the weighted average value and weighted standard deviations are calculated, where the weight is equal to number of points used for τd determination for given vesicle.

**Figure 1.**
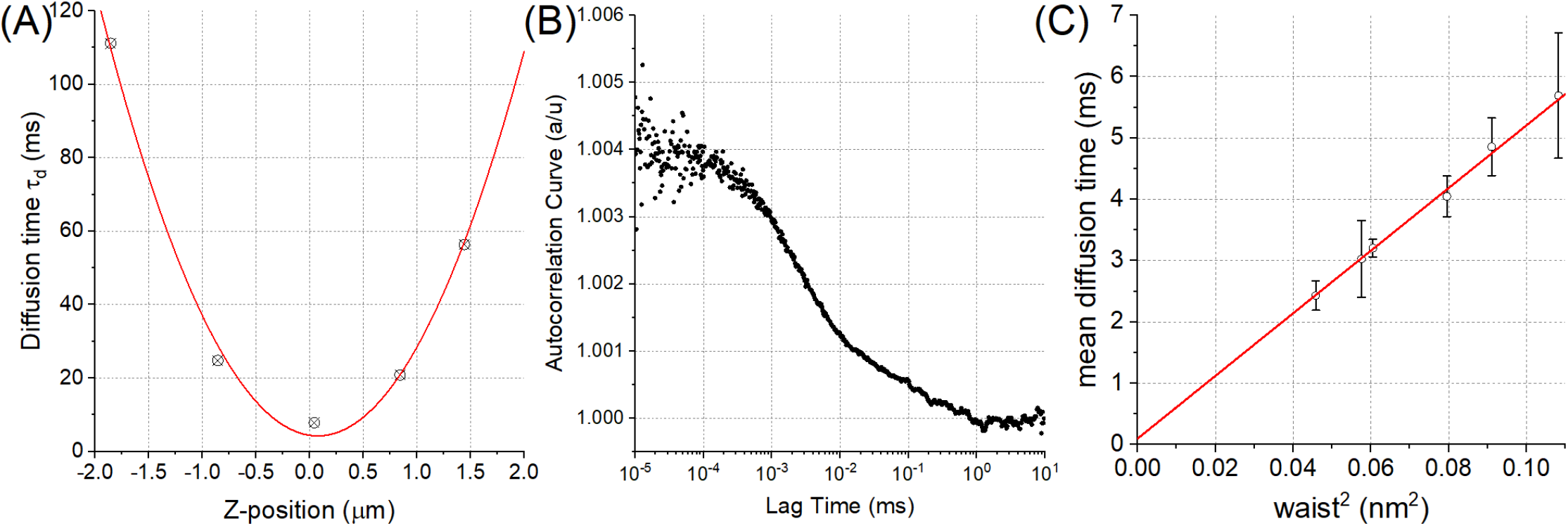
Z-scan approach combined with svFCS. (A) Determination of minimal diffusion time T_d_ for given POPC vesicle. (B) Autocorrelation function of one of such points. C) Diffusion law obtained for POPC GUVs with 1:1000 NBD-Chol.

Using this approach diffusion law for POPC GUVs presented in Figure 1.C was obtained. Interestingly, t_0_ equaled to (0,10±0,12) ms, which means that lateral diffusion in this case remains unconstrained and determined D_eff_ equaled to (4,9±0,2) um^2^/s, which should be convergent to D obtained in other classical experiments. Indeed, reported results are only slightly higher, D=8.87 μm^2^/s in 298K [18], which can be explained by lower temperature of our system. Consequently, we show that POPC bilayer is homogeneous and svFCS can be used to investigate the existence of domains on the lipid membrane systems.

To this end, diffusion law was established for POPC:C1P18:1 GUVs with NBD-Chol (Figure 2.A). Obtained value of D_eff_ equaled to (3,81±0,15) μm^2^/s. Interestingly, the value of t_0_ equaled to (2,36±0,16) ms. Such results suggest an existence of certain constrains within this lipid bilayer. There is no literature report about phase behavior of C1P18:1, but based on results obtained for POPA analogue [19], it is most likely that the membrane exhibits L_o_/L_d_ phase coexistence.

**Figure 2.**
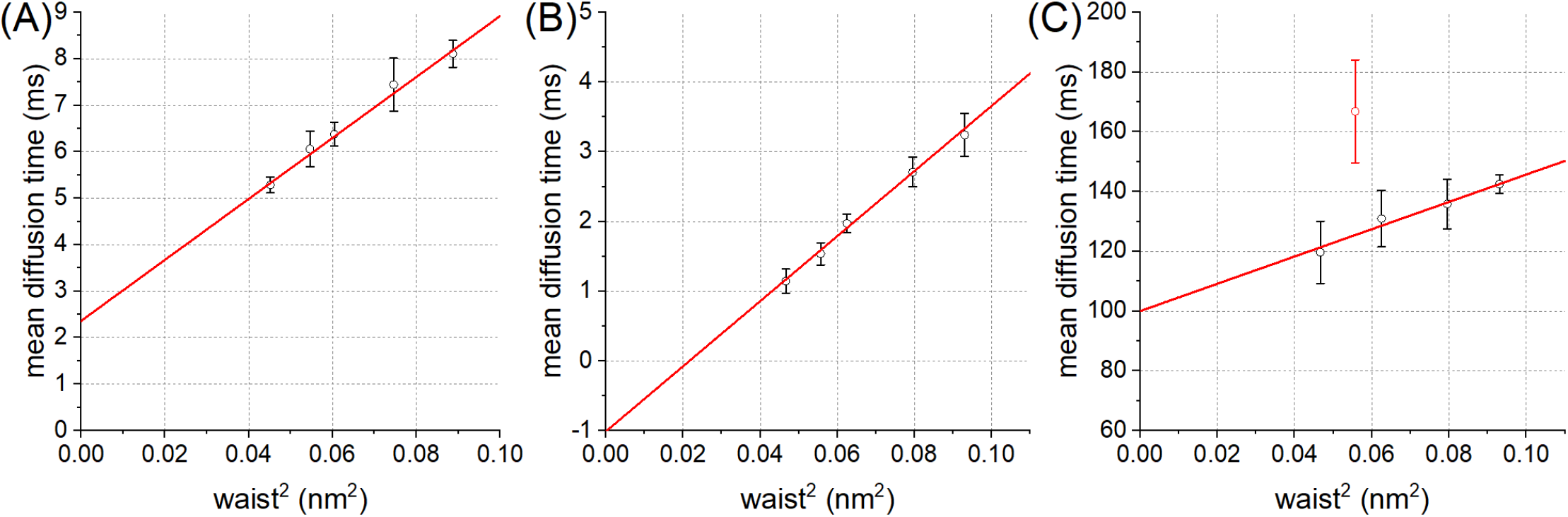
Diffusion laws obtained for GUVs containing POPC and C1P (molar ratio 8:2) with 0,1mol% NBD-Chol (A) C1P18:1, (B) first population C1P16:0, (C) second population in C1P16:0.

The most interesting results were obtained for GUVs with saturated species of C1P. C1P16:0 was reported to induce gel domains [15, 20]. Fluorescent modification of C1P16:0 was also reported to create more rigid and ordered membrane structure [21]. Contrary to previous case, even the shape of autocorrelation curves suggests two populations - most likely the fluid and gel phases. Such effect of phase on diffusion law was already reported for DMPC [22]. Hence, the obtained data had to be analyzed separately for first and second population. The first, faster, population resulted in diffusion law presented in Figure 2.B. Obtained value of D_eff_ was equal to (5,35±0,28) um^2^/s, which is similar to homogeneous POPC vesicles. Interestingly, also negligible lipid movement constraint was observed, which resulted in t_0_ equal to (−1,02±0,16) ms. Negative t_0_ correspond to case of adjacent domains separated by barriers. Most likely this constraint is observed due to existence of second population that serve as such a barriers and influence the lipid movement in first population. Negative t_0_ could also indicate non-spherical barriers. Especially that it was reported that variations in plasma membrane topography can influence the observed diffusion time [23].

The data for second, slower, population also allowed for plotting diffusion law, which is presented in Figure 2.C. Obtained values of D_eff_ equaled to (0,55±0,05) μm^2^/s. As in gel domains there is, generally, much slower lipid motion [22], this suggest that this population rather reflects gel phase. Obtained value of t_0_ is extremely high, it equaled (100±4) ms. This suggest strong constraint. On the other hand, contrary to reports on DMPC [22], the t_0_ value is positive for POPC:C1P16 membranes. This indicates, that observed effect is not only an effect of simple phase difference, but also negative curvature might influence the behavior of lipids in membranes.

In silico molecular dynamics studies of diffusion coefficient were performed to shed some light on diffusion coefficient s obtained experimentally. Agreement of diffusion coefficients should serve as a confirmation that the membrane organization is similar. As presented in Figure 3.A, in the case of POPC, the *in silico* value of diffusion coefficients equals (4.8±0,6) μm^2^/s [24]. This results is in agreement with D_eff_ from svzFCS. Similarly, for POPC:C1P18:1 mixture there was an agreement between diffusion coefficient values, as *in silico* we obtained (3,65±0,04) μm^2^/s. Given the fact that t_0_ value nearly zero, it is not surprising that both values (the D and D_eff_) are similar. On the other hand, in the case of POPC:C1P16:0 there was no agreement between simulation and experimental diffusion coefficients. The simulated one equals to (2,33±0,04) μm^2^/s. Remarkably, the simulation of single component C1P16:0 membrane resulted in D=(0,55±0,02) μm^2^/s (see Figure 3.B). Taken together, it could be stated that the two populations identified by svzFCS closely refer to diffusion coefficient to single-component C1P16:0 membrane and to single-component POPC system. This further supports that C1P16:0 form gel domains which coexist with fluid phase within the POPC/C1P16:0 vesicles and our experimental setup is able to detect such features of the studied membranes.

**Figure 3.**
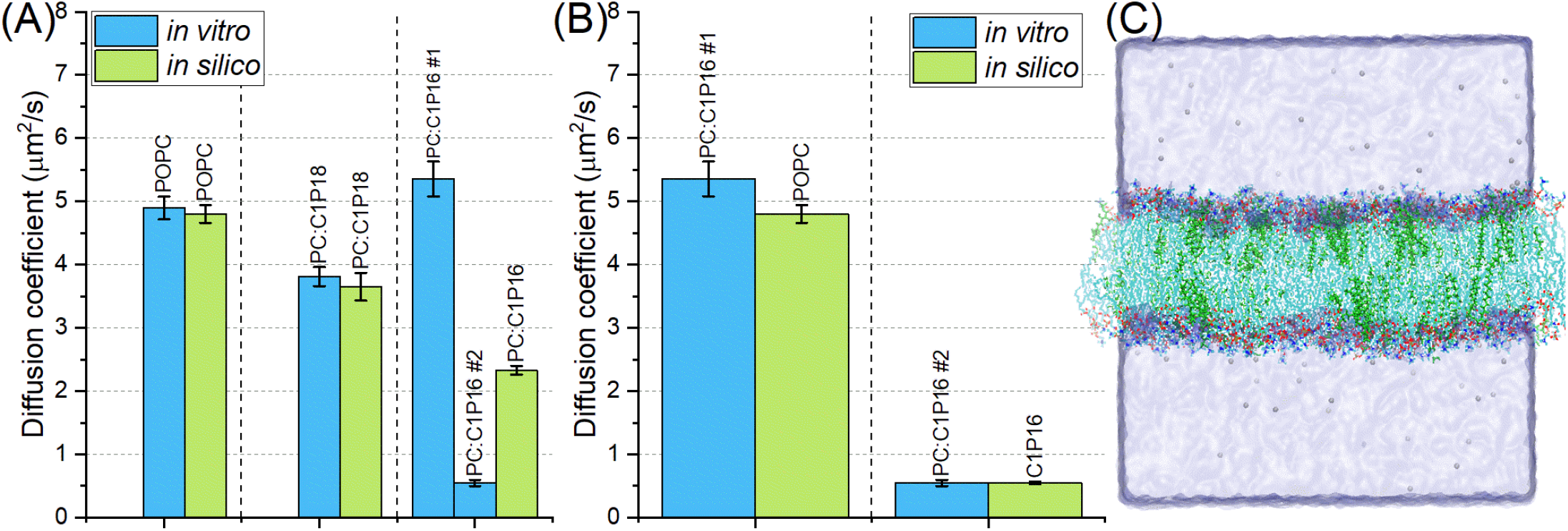
(A) Comparison between D_eff_ obtained from svFCS and respective *in silico* diffusion coefficient values obtained from Molecular Dynamics. (B) Comparison of C1P16:0 different species from svFCS to other *in silico* membrane systems. (C) Snapshot of POPC:C1P16 system from molecular dynamics studies. C1P16 molecule is colored green.

In addition to described above nano-domains, both C1P molecules were also able to form larger domains within GUVs observable via confocal microscopy as areas with non-detectable fluorescent lipid analogs. To ensure, that the observed separation is not a result of probe-induced artefacts, three probes were tested (Atto488-DOPE, Fast DiO and NBD-Chol). In each case phase separation occurred for both vesicles with C1P16:0 and C1P18:1. The domains observed on vesicles with C1P16 were edgy and not spherical, as presented in Figure 4., which is related to their gel nature. The average area of the domains for all of recorded vesicles was equal to (1,94±3,11) μm^2^, although largest domains were up to 15 μm^2^. This means that (5,11±3,63)% of the whole vesicle area were occupied by these domains.

**Figure 4.**
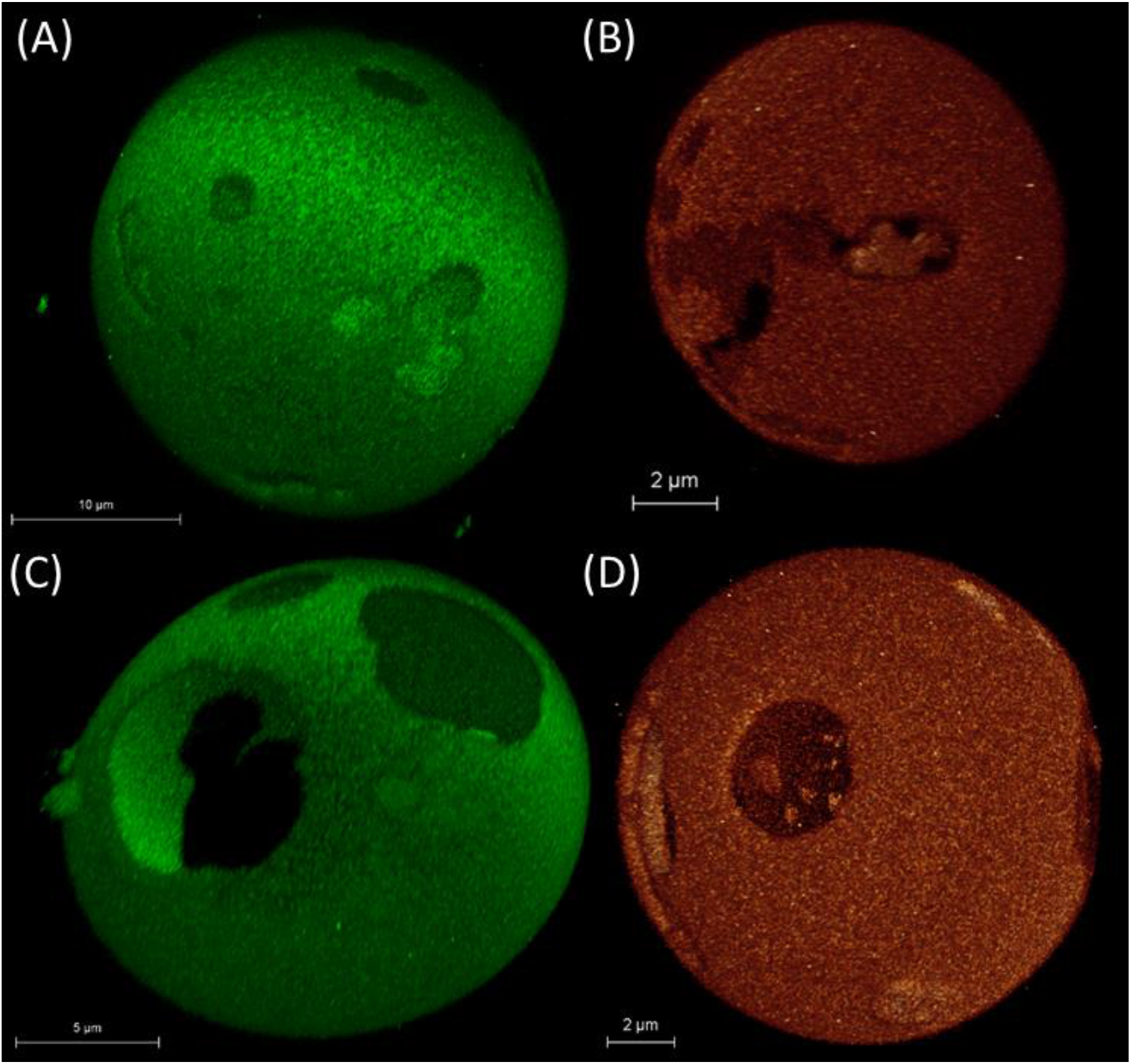
Confocal microscopy Z-stack 3D visualizations of GUVs from POPC:C1P16 8:2 mixture labelled with (A) Atto488-DOPE, (B) fast DiO and POPC:C1P18 8:2 mixture labelled with (C) Atto488-DOPE and (D) fast DiO.

In the case of vesicles with C1P18:1 the domains were round and spherical (as presented in Figure G.B) which suggest that they are fluid/liquid. Their average area was significantly larger than those observed for C1P16 and was equal to (20,1±18,8)μm^2^ for all of recorded vesicles, while largest domains reached up to 80 μm^2^. On average they cover (13,0±2,4)% of vesicle area.

In summary, we’ve established a new methodological approach for the measurement of diffusion law in GUVs. It was used to determine diffusion parameters of vesicles containing C1P. In case of control POPC membrane obtained value of t_0_ was practically equal to zero, nicely reflecting lack of any constrains or coexistence of domains. More interestingly, for both POPC:C1P16:0 and POPC:C1P18:1 bilayers the values of t_0_ were not equal to zero, which suggests an existence of some kind of constraint and most probably existence of nano-domains. In the case of membrane with C1P16:0, clearly two populations were even visible in autocorrelation curves. Comparison with diffusion coefficients obtained *in silico* suggest that one represents nano-domains overwhelmingly occupied with C1P, while the other reflects POPC-enriched regions. In the case of C1P18:1 only one population was observed that was in strong agreement with *in silico* results. These conclusions also correlate with the character of domains microscopically observed on GUVs. Deciphering behavior of various species of C1P, one of the key signaling lipids in cells, is of vital importance to understand a variety of molecular mechanisms governed by this lipid. Domains of various character formed by C1P may be important for efficient fine-tuning recruitment and/or activity of proteins involved in cell-signaling pathways.

## Experimental and Computational methods

GUVs are obtained using electroformation method in sucrose solution (120mOsm). Agarose immobilization [25] of vesicles (up to 1% w/v) was used to remove issue of vesicle movement during the measurement. Measurements on both confocal microscope and svFCS were done in 20°C. Molecular dynamics studies were performed using NAMD software [26] with CHARMM36 [27] force fields under NPT conditions for at least 30 ns. Diffusion coefficient was established with Diffusion Coefficient Tool plugin [28]. More details about details of experimental setup and analysis and computational studies are presented in Supplementary Information.

## Acknowledgements

This work was possible thanks to the financial support from the National Science Centre (Poland) grant no 2018/30/E/NZ1/00099. The authors would like to thank dr Tomasz Trombik (current address: Medical University of Lublin) and Karolina Wójtowicz (University of Wrocław) for their support with svFCS approach. We would also like to thank dr hab. Sebastian Kraszewski for valuable input in C1P force field modification.

