## Supporting Information for "Investigation of nano- and micro-domains formed by ceramide-1-phosphate in freestanding lipid bilayers"

### 1 Experimental details

#### 1.1 Materials

Lipids POPC (1-palmitoyl-2-oleoyl-glycero-3-phosphocholine), C1P 18:1/d18:1 (N-oleoyl-ceramide-1-phosphate) and C1P 16:0/d18:1 (N-palmitoyl-ceramide-1-phosphate) were purchased from Avanti Polar Lipids (Alabaster, AL, USA). Fluorophores NBD-Cholesterol was purchased from Avanti Polar Lipids (Alabaster, AL, USA), fast-DiO was purchased from ThermoFisher Scientific (Massachusetts, USA) and Atto488-DOPE was purchased from Sigma Aldrich (Missouri, USA). Structures of lipids are presented in Fig S1. Sucrose and glucose were purchased from Sigma Aldrich (Missouri, USA). Fisher Scientific Low-melting temperature agarose polymer ($T_m$~65°C, $T_g$~25°C) was purchased from Fisher Scientific (Massachusetts, USA). The ultra-pure water used in experiments was obtained from water purification system (Milipore).

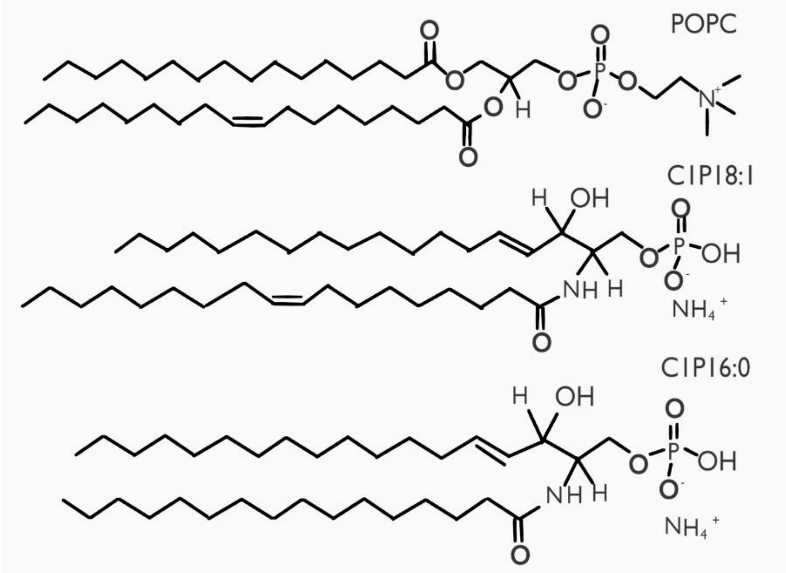


Figure S1. Structures of lipids used in the study.

#### 1.2 GUVs electroformation and agarose immobilization

The modified method of model membrane formation for giant unilamellar vesicles (GUV) was used. Briefly, 10 µl of 1 mM POPC/C1P mixture and fluorescent probe mixture (0.1mol% for svzFCS, 0.5mol% for confocal visualization) in chloroform was distributed equally along the platinum electrodes and dried under the vacuum for 1 hour. The electrodes were then submerged in aqueous non-conductive sucrose solution (120mOsm) and a square 10 Hz AC electric field was applied for 10 h with 1V voltage in a custom PTFE (polytetrafluoroethylene) electro-formation chamber increasing with 1V every hour up to 4V [1].

Agarose immobilization was done according to protocol described by Lira *et al* [2]. Briefly, agarose was dissolved in 120 mOsm Glucose solution to obtain either 1 or 2% w/v concentration depending on application. Vesicles and agarose were mixed equally in volume while the polymer was still in the fluid state (around 35–40 °C). Final 1% w/v agarose concentration was sufficient for vesicles remaining immobilized for the measurement time in svzFCS, while for visualization 0.5% w/v was applied.

#### 1.3 spot variation Z-scan Fluorescence Correlation Spectroscopy (svzFCS)

The svzFCS measurements were performed using a custom-made svFCS optical system based on a classical Axiovert 200 M fluorescence microscope (Carl Zeiss, Oberkochen, Germany). The protocol was partially based according to the one described by Mailfert *et al* [3, 4]. However, due to changing investigated sample from cells to GUVs, z-scan approach was implemented to the protocol. The z-scan FCS approach was based on results and conclusions presented by Heinemann *et al* [5]. The waist size was calibrated with 2 nM Rhodamine 6G solution and 488 nm laser beam illumination at the intensity of 330 µW. GUVs analyses were performed at 20°C while the laser beam was adjusted to 2–4 µW. After the vesicle was detected, the point of measurement was set to centre of the vesicle, which was followed by z-scan to determine the z-position of upper membrane. There were at least five z-positions for each vesicle (one in the highest intensity z-position, two above and two below). The signal was collected by a series of 6 runs lasting for 10 s each for each of z-position of the vesicle. The measurements were carried out on 5 to 8 individual GUVs and the obtained data were analysed by the IGOR Pro software (WaveMetrics). The collected autocorrelation functions were fitted with a 2D lateral diffusion model [6] (either one or two population depending on the vesicle type) and the mean diffusion time $\tau_d$ was calculated. Five to six waists were analysed in order to construct a single diffusion law. Basic visualization and calculations were performed in Excel (Microsoft Corporation)

#### 1.4 Confocal visualization studies

3D z-scans of GUVs were recorded using Stellaris 8 confocal microscope (Leica, Wetzlar, Germany) equipped with an HC PL APO 86x/1.20 water immersion objective (Leica,Wetzlar, Germany). 521×521 pixels images were recorded with a hybrid (HyD) detector and 0.5 Airy unit pinhole size. For all investigated fluorophores 488 nm incident wavelength was used. Analysis of domains is performed using custom script in MATLAB. Briefly, the centre position and radius of vesicle is determined. This is followed by determination of artificial points equally distributed on the sphere with determined radius (to ensure equal distribution of the points in pole region). For each point a rectangular slice is made from sphere centre to the point. The number of points in the vicinity of radius is calculated and, based on general quantities accustomed to given vesicles, the presence of fluorescence signal is determined.

#### 1.5 Statistics

In order to test the significance of difference between the parameters, unless specified otherwise, the one-way ANOVA test was used with the significance level at 0.05. The Tukey test was used as a post hoc test. All statistical analysis was performed using the OriginPro 2015 (OriginLabs) software. Average values are presented with standard deviation.

### 2 Computational Details

#### 2.1 Force field modification

C1P molecules force fields are not available in Charmm, hence they were created by adjusting the energies for lipid head on sphingolipid backbone. C16:0 and C18:1 Ceramide-Phosphates topologies were constructed based on CHARMM36 topologies. Specifically, sphingolipids PSM and OSM were used to reflect acyl chain region. The phosphate head region was based on both PA and DOPP2 (for double oxygen bond). Partial charges were readapted to FFCharmm consistency according to the scheme shown in shown in Fig S2.

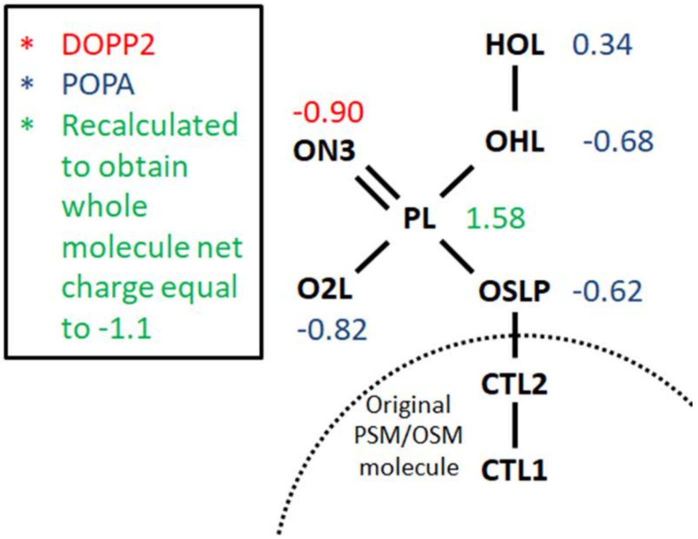


Figure S2. Detailed charge distribution of the head region of modified C1P molecule force field.

#### 2.2 Molecular Dynamics Simulations

The full-atomistic MD simulation was performed using NAMD 2.13 [7] software with CHARMM36 force fields [8] under NPT conditions (constant: Number of particles, Pressure and Temperature). Lipid membrane systems consisted of 648 lipid molecules (324 on each of leaflets). System was hydrated in such a way that 75 water molecules per lipid molecule was used. All systems were additionally neutralized with positive counter ions. All systems were equilibrated using standard equilibration procedure. Total simulation time for all systems was at least 30 ns with last 10 ns used for analysis. Simulations were carried under 22°C (285.15K). The Diffusion Coefficient Tool plugin was used for lipid molecules lateral (2D) diffusion determination [9]. It was calculated using Einstein's relation with mean square displacement (MSD) of phosphorus atoms of the chosen molecular species.

### 3. Z-fits in sv-zFCS

In this section examples of Z-fits for sv-zFCS data was presented along with corresponding autocorrelation curves. The results are presented for POPC (Fig S3), POPC:C1P16:0 (Fig S4) and POPC:C1P18:1 (Fig S5).

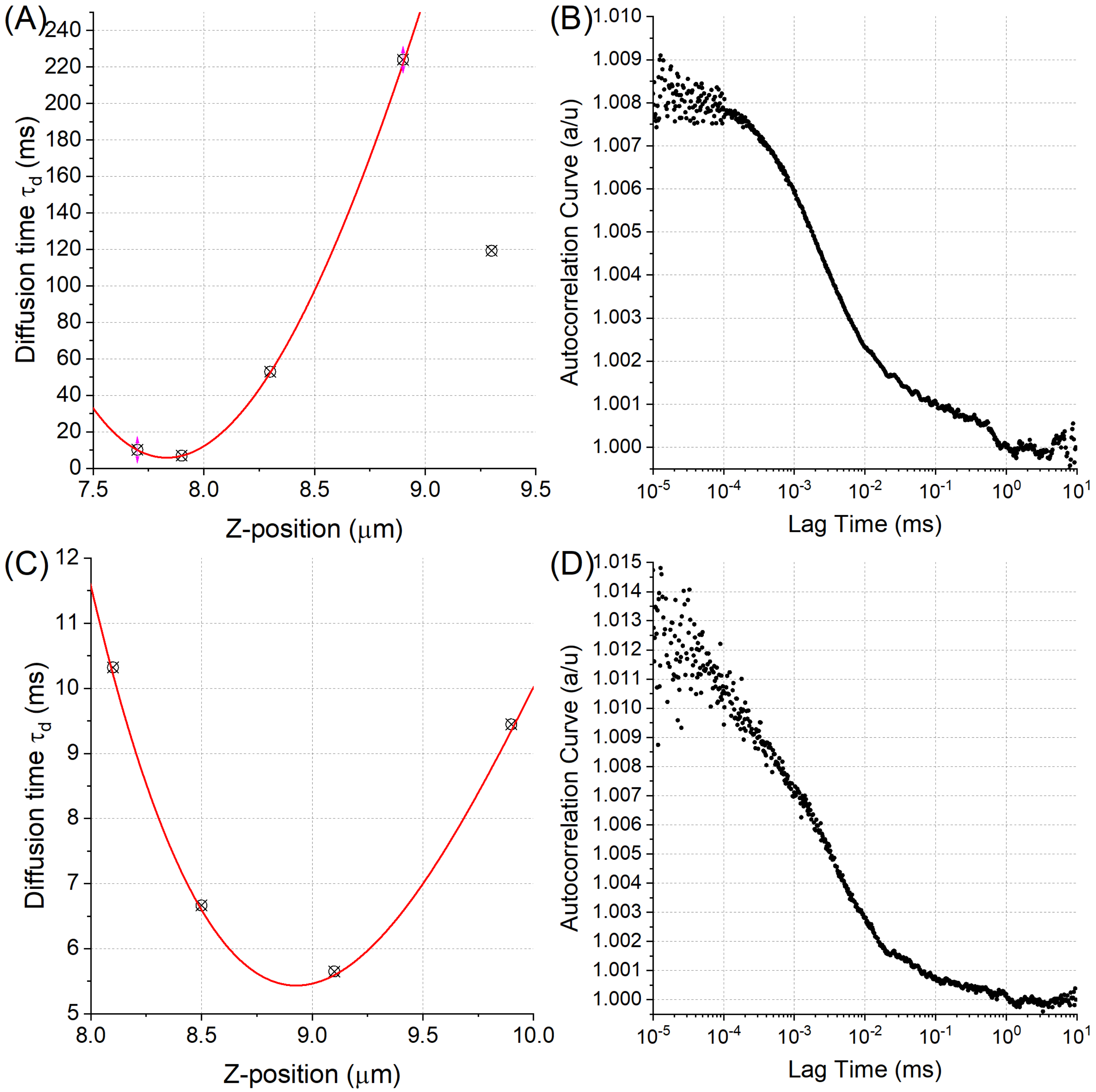


Fig S3. Z-scans and corresponding autocorrelation curves for POPC vesicles with waist set to (A-B) 236 nm and (C-D) 282 nm.

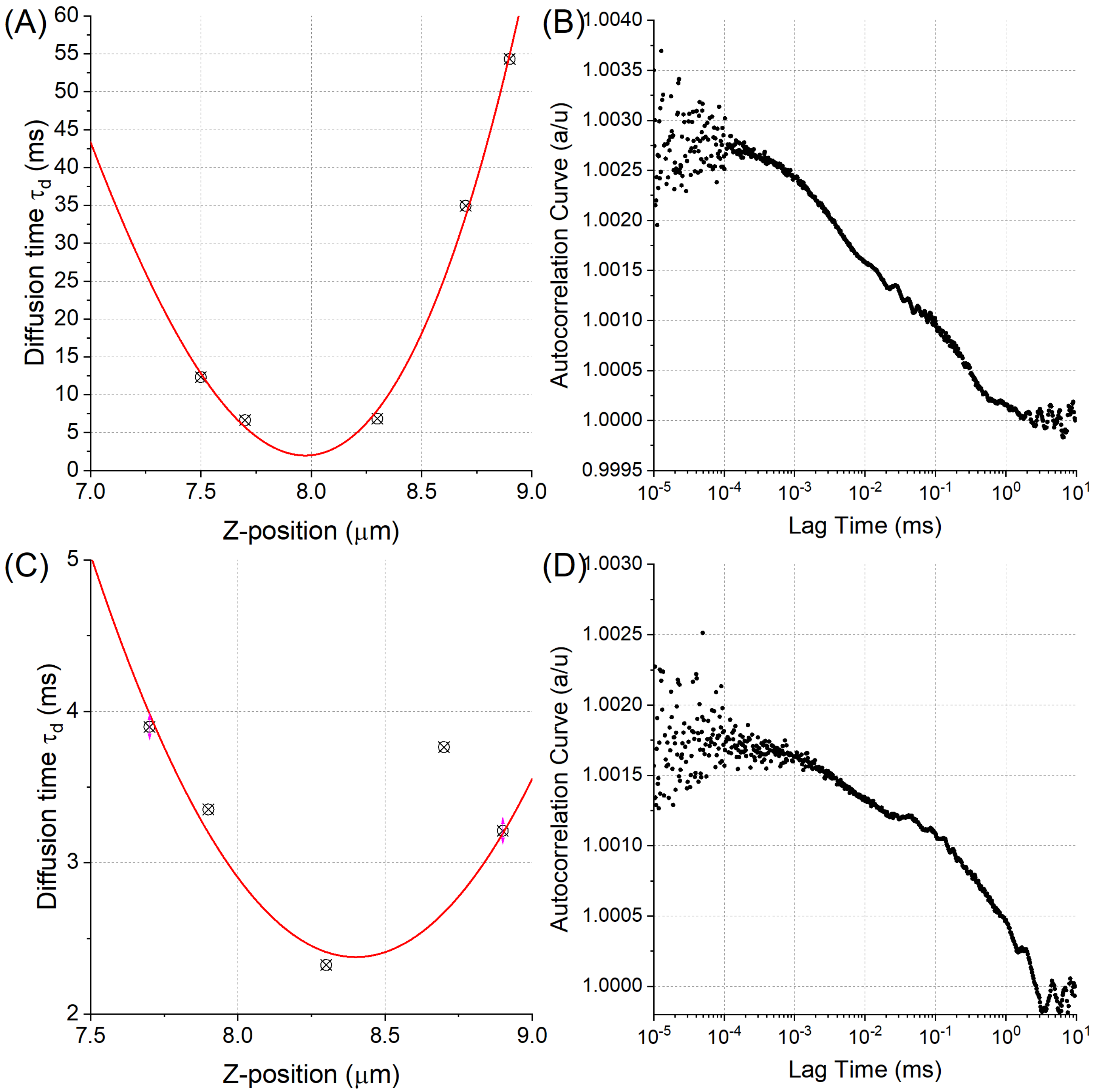


Figure S4. Z-scans and corresponding autocorrelation curves for POPC:C1P16:0 vesicles with waist set to (A-B) 216nm and (C-D) 305nm.

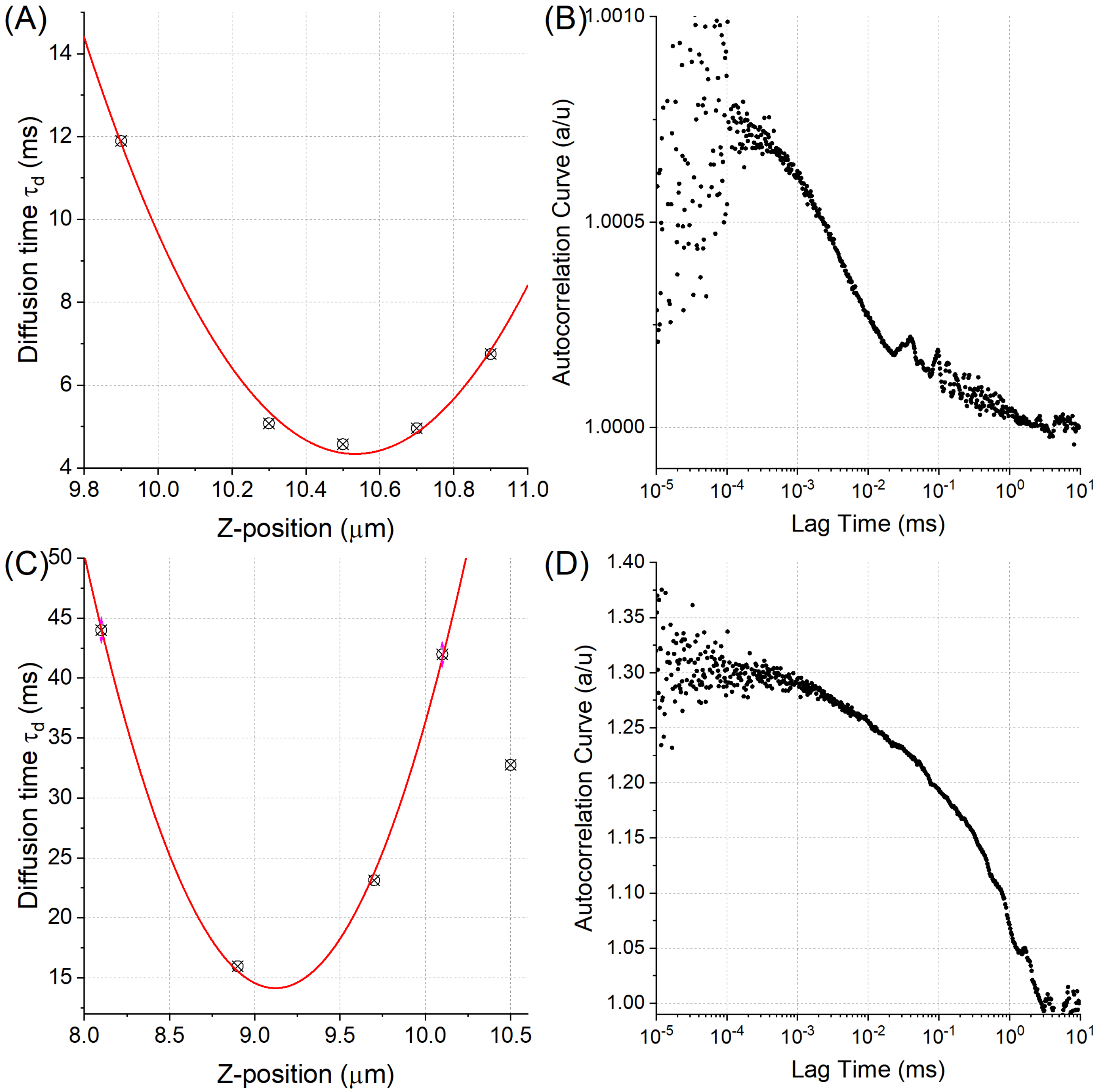


Figure S5. Z-scans and corresponding autocorrelation curves for POPC:C1P18:1 vesicles with waist set to (A-B) 212nm and (C-D) 273 nm.

**4. 3D reconstructions of giant unilamellar vesicles**

In this section the 3D reconstructions for vesicles with POPC:C1P18:1 (Fig S6 A-I) and POPC:C1P16:0 (Fig S7 A-I) are presented. Vesicles were labelled with Atto488-DOPE, DiO and NBD-Chol. At least 3 vesicles are shown for each of the dye.

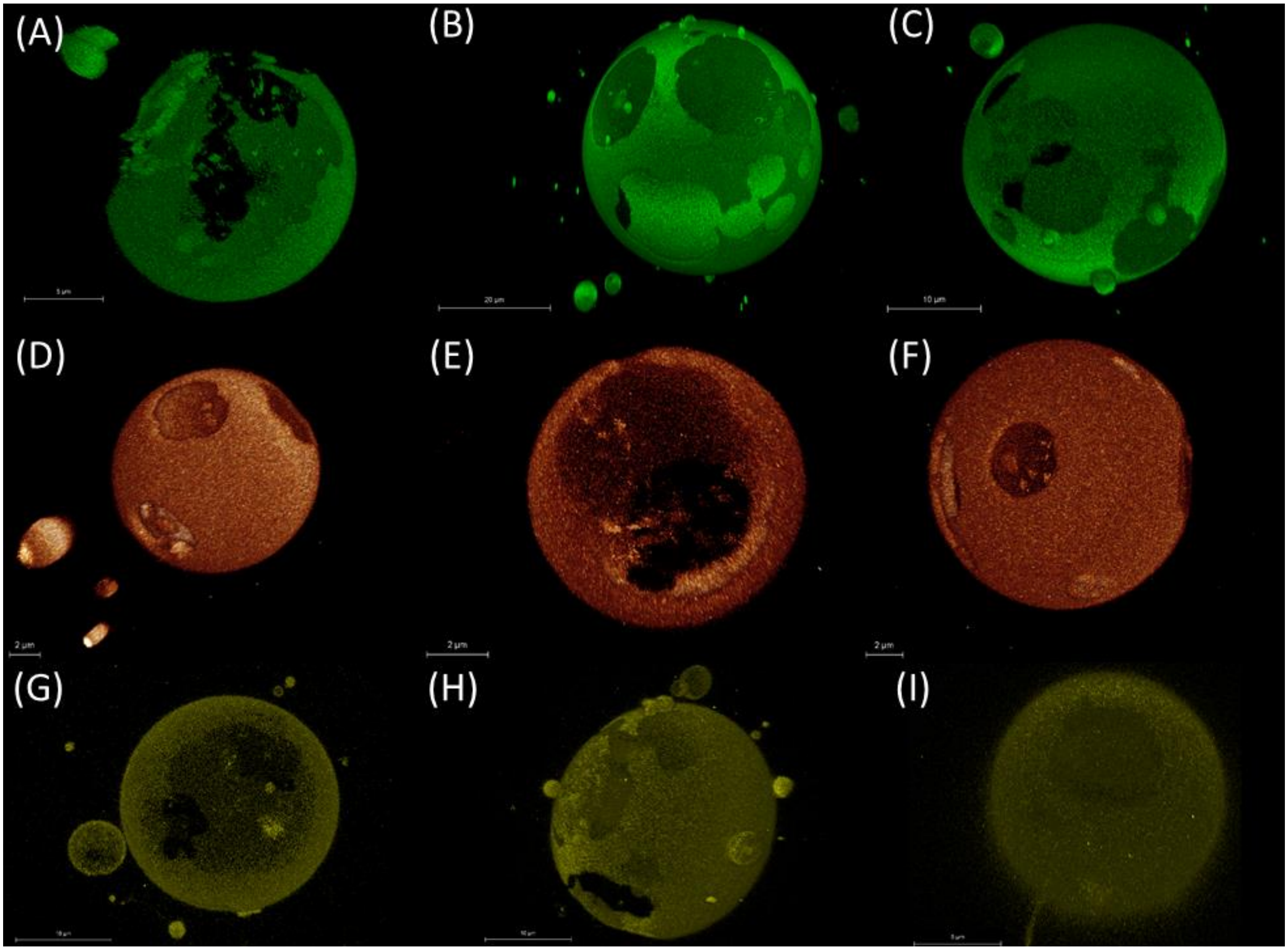

Figure S5. 3D reconstructions of POPC:C1P18:1 labelled with (A-C) Atto488, (D-F) DiO and (G-I) NBD-Chol.

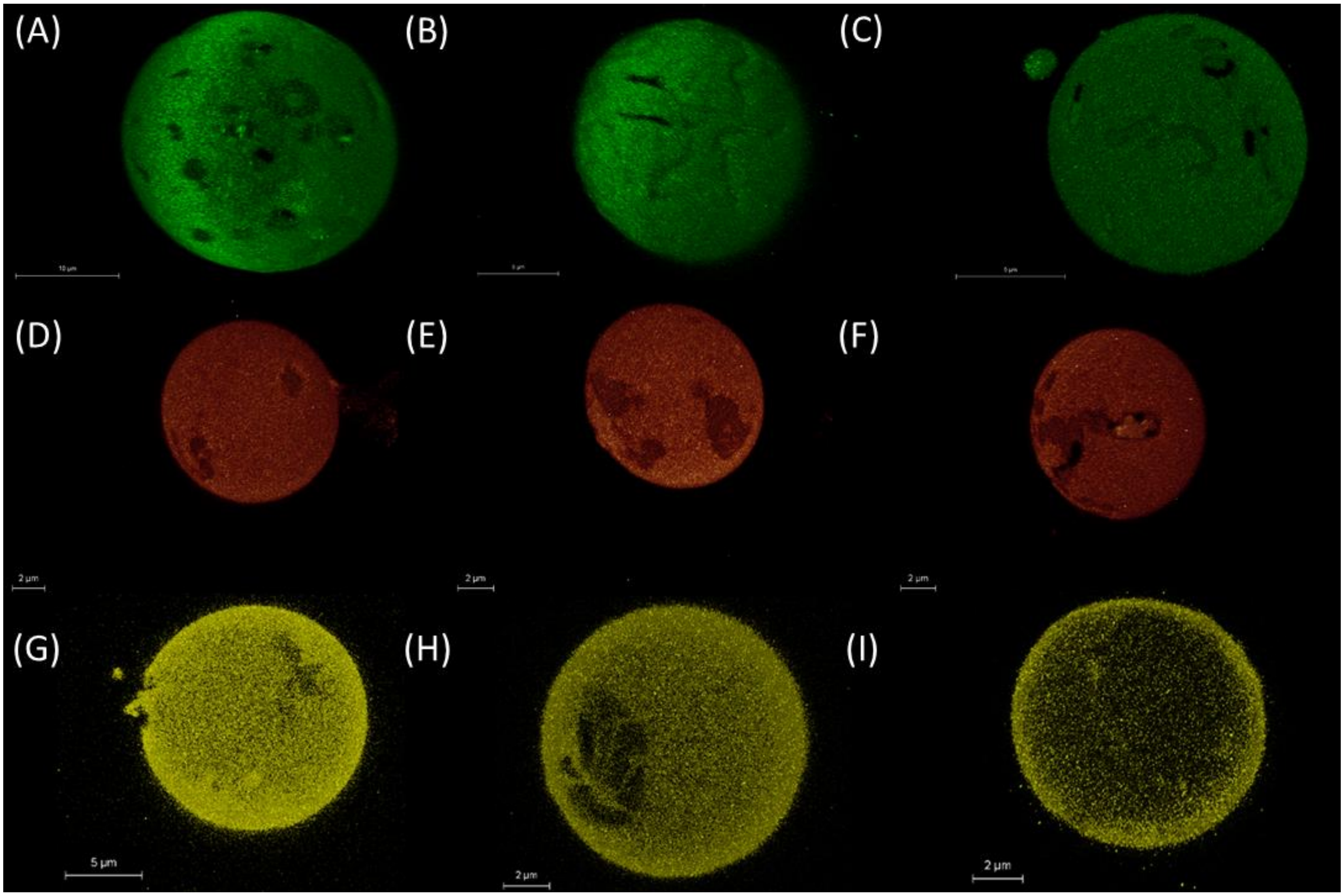

Figure S7. 3D reconstructions of POPC:C1P16:0 labelled with (A-C) Atto488, (D-F) DiO and (G-I) NBD-Chol.
